# Promoting oak (*Quercus pubescens* Willd.) seedling development: interactions between canopy opening and competition with grass (*Festuca ovina* L.) under soil water deficit

**DOI:** 10.64898/2026.02.04.703752

**Authors:** Solène Brasseur, Mathieu Santonja, Tanguy Sanet, Christine Ballini, Sylvie Dupouyet, Bernard Prévosto, Anne Bousquet-Mélou

## Abstract

To adapt Mediterranean forests to increasingly harsh climatic conditions by promoting genetic diversity, thinning is often considered an effective strategy to enhance sexual regeneration within population diversity. However, determining an optimal thinning level that both increases local irradiance and maintains favorable microclimatic conditions for germination, without excessively promoting herbaceous competition, remains challenging. To better understand how abiotic and biotic factors influence oak seedling development and to help identify a balanced thinning level under climate change, we conducted a semi-controlled experiment testing the combined effects of competition with a grass species (Poaceae), two canopy opening levels, and soil water deficit. Our results highlight the crucial role of competition with Poaceae species - in our case *Festuca ovina* L. - in oak regeneration. Their presence not only intensifies competition for essential resources, but also modifies soil properties and alters belowground interactions, overall creating less favorable conditions for oak seedling establishment. In addition, our results highlight the significant impact of key abiotic factors like canopy opening that influences local irradiance and soil water deficit, as well as their interactions with the effects of competition. We observed a consistent need for adequate local irradiance to ensure optimal seedling performance, suggesting that successful regeneration depends on balancing sufficient canopy opening to improve light availability with maintaining sufficient cover to mitigate water stress and limit grass competition. Overall, our study contributes to the broader debate on sustainable forest management strategies under changing climatic conditions.

## 1. Introduction

Species from the genus *Quercus* have often been favored in Europe by foresters due to the economic value of oaks (Marchi et al., 2016; Timbal & Aussenac, 1996). A large part of the oak forests has been traditionally managed as coppices, supplying firewood and wood for charcoal production since centuries (Marchi et al., 2016; Nocentini & Coll, 2013; Picchio et al., 2011). This ancient and simple management method relies on vegetative regeneration with harvest rotations generally ranging from 15 to 35 years depending on the tree species, growth conditions and desired products (Bottalico et al., 2014; Chirici et al., 2020; Spinelli et al., 2016). Stump sprouting and rapid growth after felling (mainly clearcutting) allows shorter harvest intervals than in high forests that are regenerated by seeds (Ducrey & Turrel, 1992; Giovannini et al., 1992).

Oak coppices are widespread worldwide and especially common in Mediterranean regions in France, Italy or Spain (Fabbio, 2016; Marchi et al., 2016). However, after the Second World War, coppicing was largely abandoned (Prévosto et al., 2013; Salomón et al., 2017; Vadell et al., 2022), leading to severe within-coppice competition among shoots on unmanaged stools, driving standing stands toward growth stagnation and structural collapse (Cañellas et al., 2004). Furthermore, over the long term, this lack of management implied a significant reduction of the vegetative regeneration due to reduced sprouting efficiency in abandoned and ageing coppices (Ladier et al., 2014; Prévosto et al., 2013). This limitation on regeneration poses a major challenge to forest renewal, at a time when biomass has returned to the forefront as a potential energy source (Prévosto et al., 2013; Santi et al., 2024).

Mediterranean forests will face increasing water deficit combined to rising temperatures due to ongoing climate change (IPCC, 2023; Lionello et al., 2008, 2014). These unfavorable abiotic conditions for tree growth are often combined with an increase in insect outbreaks or fungal attacks, affecting forest health and potentially leading to dieback, defoliation and even mortality (Allen et al., 2015; Lemaire et al., 2022). In this context, adaptive solutions that effectively strengthen the long-term resilience of forests, and in particular their capacity to regenerate from seeds (sexual regeneration), are needed (Vilà-Cabrera et al., 2018). Sexual regeneration implies genetic mixing and environmental selection of seedlings, ensuring better adaptation to the environment (Bussotti et al., 2015; Vilà-Cabrera et al., 2018). However, particularly since the beginning of the 21^st^ century, this regeneration pathway has faced significant challenges (Plieninger et al., 2010; and Tyler et al., 2006 for Californian oaks), including mainly insufficient recruitment, slow-growing seedlings and low survival rates (Gauquelin et al., 2016; Prévosto, 2020a).

To adapt forest systems to these new and harsher climatic conditions, thinning is often considered an effective method to enhance tree growth, improve their health status and favor seed regeneration by reducing competition for resources between remaining trees (Kerr & Haufe, 2011) and limiting the impact of drought on growth (Cabon et al., 2018; Sohn et al., 2016). Thinning is also an adaptive solution that can contribute to soil carbon sequestration (Mäkipää et al., 2023; Zhang et al., 2018), biodiversity provisioning (Li et al. 2020) and fire risk reduction (Taylor et al., 2021). It also stimulates fruit-production (number of seeds per tree) (Healy et al. 1999; Rodriguez-Calcerrada et al., 2011). Furthermore, compared to coppicing, this management method is considered an adaptive solution for promoting sexual regeneration by enhancing seed production (Otto et al., 2012) and favoring long-term establishment of seedlings by promoting their survival and growth (Bran et al., 1990).

Seedling development is influenced by numerous ecological filters, such as soil properties and microclimatic conditions (*e.g*., soil moisture, local irradiance or temperature fluctuations) (Mendoza et al. 2009). Both abiotic and biotic factors influence seedling establishment and growth, making it difficult to determine a balanced thinning level (Mölder et al., 2019). Light and water availability for instance, play a key role but are influenced by the intensity of thinning (Prévosto et al. 2011; Retana et al., 2011). First, seedling growth benefits from increased local irradiance due to opening of the canopy (Lusk et al., 2008; Niinemets, 2006; Quero et al., 2007). However, the canopy is known to also produce favorable microclimatic conditions for seed germination (Mölder et al., 2019; Retana et al., 1999), with the forest cover acting as a buffer for extreme temperatures and high vapor pressure deficit (Prévosto, 2020a; Valladares et al., 2016). Moreover, the positive effect of opening on seedling growth could be offset by reduced water availability and competition for soil nutrients with the herbaceous stratum that is promoted by forest opening (Gaudio et al., 2011; Wagner et al., 2010). In particular, grass species of the Poaceae family are known to produce a dense root system with a high ability to uptake water and nutrients (Gibson & Newman, 2001; Yamada, 2011), and are recognized as strong competitors to tree seedlings (Balandier et al., 2006; Schaller et al., 2003; Vernay et al., 2018). Competition faced by young tree seedlings can also be mitigated by the ability of grass species to influence the seedlings’ ability to access non-limiting resources. The negative effects of Poaceae species on the root systems of young tree seedlings or on mycorrhizal symbiosis (which usually promote their establishment, growth and productivity), for example, may affect their ability to absorb water and nutrients (Fernandez et al. 2023). For example, young *Quercus petraea* (Matt.) Liebl. seedlings competing with *Avenella flexuosa* (L.) Drejer (formerly *Deschampsia flexuosa* (L.) Trin.) were shorter in height, had fewer branches and a smaller root system (Collet et al., 1996). Similarly, *Molinia caerulea* (L.) Moench restricted both *Q. petraea* root growth and ectomycorrhization rate (Fernandez et al., 2023). Finally, although the competitive role of the herbaceous layer is often emphasized, facilitative effects have also been reported. For example, Tonioli et al. (2001), in a field experiment, demonstrated that surrounding vegetation, while competing with young *Quercus pubescens* Willd. seedlings for resources, may also facilitate seedling emergence.

*Quercus pubescens* Willd., commonly called downy oak, is native from the Mediterranean basin and characteristic of the supramediterranean bioclimate (Quézel, 1979) and is found from southern Europe to southwestern Asia (Ganatsas & Tsakaldimi, 2013), on hillsides generally between 200 and 800 m a.s.l. (Pasta et al. 2016) and up to 1400 m a.s.l. in France (Rameau et al., 2008). This deciduous species exhibits some tolerance to moderate summer drought and low winter temperatures, but is vulnerable to extreme climatic events, including recurrent frost and intense droughts, which are characteristic of continental climates (Pasta et al., 2016). It grows on both acidic and calcareous soils, though it shows a marked preference for the latter, and is characterized by a heliophilous and thermophilous ecological strategy (Contran et al., 2013; Pasta et al., 2016). In France, *Q. pubescens* is the dominant species on approximately 1.41 million hectares, of which 841,000 hectares are forests where it comprises over 75% of the canopy cover (IGN, 2024). In south-eastern France, these forests are mainly managed as coppices. However, the cutting frequency has decreased over the past few decades, and many *Q. pubescens* stands are now left without management and ageing (Ladier et al. 2014).

To better understand how abiotic and biotic factors influence the development of *Q. pubescens* seedlings and help identify a balanced thinning level in a climate change context, we conducted an experiment under semi-controlled conditions. We tested the effects of competition with a Poaceae species (*Festuca ovina* L.), combined with two levels of simulated canopy opening and soil water deficit. We analyzed the effects of these factors on biomass allocation to the different plant compartments, as well as on some key morphological and physiological variables in young *Q. pubescens* seedlings. As competition with grass species may influence the nutritional status and water supply of oak seedlings, we also examined the seedling root mycorrhization and soil physico-chemical characteristics. We focused on the interactions between these three factors (*i.e.*, canopy opening, soil water deficit and competition) and tested the following hypotheses: 1) Seedling development, assessed through morphological, physiological, and functional parameters, is limited when forest canopy opening is low and soil water deficit is high; 2) Competition with *F. ovina* has a negative effect on seedling morphological, physiological, and functional parameters (height, diameter, aboveground and underground biomass, leaf surface, number of leaves, SLA, fungal abundance *etc.*); 3) the negative effect of competition is limited under low canopy opening level and soil water deficit.

## 2. Materials & Methods

### 2.1. Plant materials

#### Target species: *Quercus pubescens* Willd

For our experiment, we used *Q. pubescens* acorns collected at an experimental site in Saint-Christol d’Albion, SE France, 44°02’58.0’’N 5°32’35.8’’E, 875 m a.s.l., which was established as part of the H2020 HoliSoils project (https://holisoils.eu). This stand is a *Q. pubescens* mature coppice that was last harvested 70 to 80 years ago (see Brasseur et al. (2025) and Ménival et al. (2025)) and for further study site description). Saint-Christol-d’Albion is characterized by a supramediterranean bioclimate with an average annual precipitation of 980 ± 47 mm and an annual mean temperature of 10.1 ± 1.1 °C (mean ± standard error calculated for the 1995-2024 period with the Saint-Christol-d’Albion climatological station data, Météo-France 2024). Acorns’ collection was conducted while maximizing the diversity of maternal trees sampled across the 5-ha of the experimental site.

In October 2022, over 100 acorns were collected (after visual inspection and controlled with the floating method) and stored at 4°C in humid sand until germination. In January 2023, they were planted in individual 1L pots filled with a substrate composed of 50% universal peat-free potting substrate (coconut fiber, vegetable compost and bark compost mix) and 50% soil collected from the naturalized green spaces of Saint Jérôme campus of Aix Marseille University (AMU). The pots were placed for a year in the greenhouse of the IMBE experimental garden (AMU Saint Jérôme Campus, Marseille, France) (Figure 1). They were watered every 3–4 days to 100% of their field capacity.

**Figure 1.**
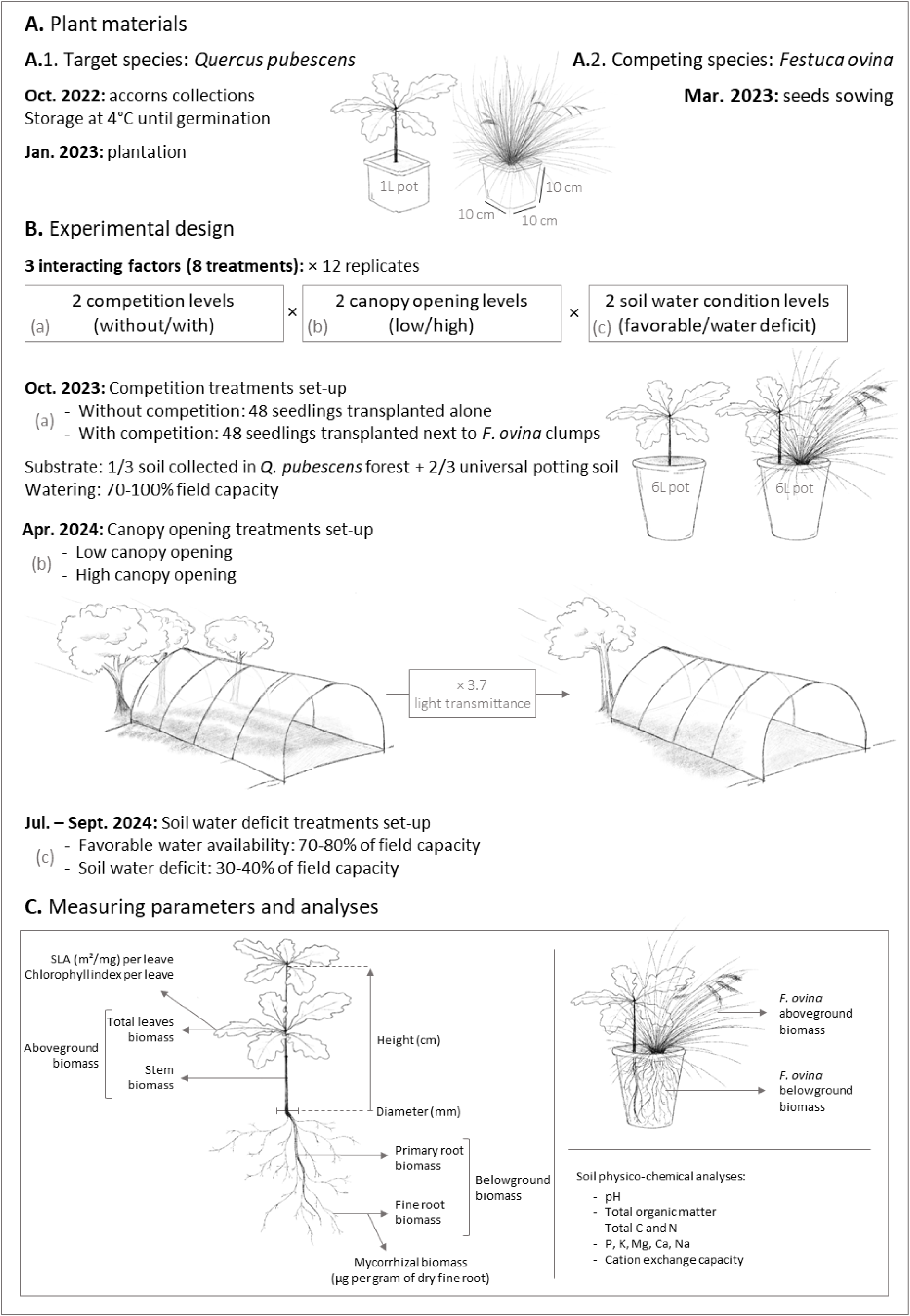
Experimental design testing 2 competition levels, 2 canopy opening level and 2 soil water condition levels on *Q. pubescens* seedling growth.

#### Competing species: *Festuca ovina* L

The genus *Festuca* L. (Poaceae) is commonly found in the meadows and steppes of North America and Eurasia (Daniel, 2001). It is a perennial plant growing in dense clumps that occasionally spreads through short rhizomes (Yamada, 2011). This genus is particularly known for its drought tolerance thanks to a fibrous and extensive root system (Gibson & Newmann, 2001; Yamada, 2011).

*Festuca rubra* L. and *Festuca glauca* L. are both present at the Saint-Christol experimental site, particularly in the most open plots. Due to the plant material accessibility, we used *Festuca ovina* L. as study species, as it shares ecological characteristics with *F. rubra.* They are both heliophilous species that grow on moderately acidic soils (Rameau et al., 1989). *Festuca ovina* seeds were supplied by Les Semences du Puy SARL (origin Denmark, harvest in 2019) and were sown in March 2023 in 10 cm deep trays filled with 2/3 universal potting substrate and 1/3 perlites and the plants were grown in the greenhouse (Figure 1).

### 2.2. Experimental design

The experimentation was set up at the IMBE experimental garden (Aix Marseille University, Saint-Jérôme Campus, Marseille, France). The climate is typically Mediterranean with a mean annual temperature of 17.1 ± 2.7 °C and mean annual precipitations of 443.7 ± 63.9 mm mainly distributed from September to January and from April to May (mean ± standard error calculated for the 2019-2024 period with the Marseille Observatory climatological station data, Météo-France (2024)).

The experimental design included three interacting factors: 2 competition levels (with/without) x 2 canopy opening levels (high/low) x 2 soil water condition levels (water deficit/favorable water condition), with 12 replicates per combination for a total of 96 pots.

*Quercus pubescens* seedlings were transplanted in 6L pots (19 × 19 cm and 21 cm deep): 48 seedlings transplanted alone in the pots (*i.e.*, without competition treatment) and 48 seedlings transplanted next to a 10 × 10 cm^2^ *F. ovina* clumps (*i.e*., with competition treatment) (Figure 1). The 96 seedlings selected exhibited highly homogenous morphological traits (similar shoot height, collar diameter, and overall development). The substrate was composed of 1/3 soil collected on the Saint-Christol-d’Albion study site and 2/3 universal potting soil (physico-chemical analysis of the substrates at the beginning of the experiment is presented in Table 1). The use of natural soil guarantees the presence in the pots of microorganisms naturally present in the *Q. pubescens* forest of Saint-Christol-d’Albion, while the use of “universal soil” ensures adequate drainage despite the relatively high clay content of the forest soil. The plants were kept under a plastic tunnel under stable watering (70-100% field capacity) and canopy opening conditions for 6 months. Soil water content was controlled by monitoring pot weights every 2-3 days.

**Table 1.** Means ± standard errors of substrate physico-chemical parameters at the beginning of the experiment (n=6).

|  |  |
|---|---|
| CEC (meq/100 g) | 20.6 ± 0.7 |
| pH (Water) | 5.4 ± 0 |
| pH (KCl) | 4.8 ± 0 |
| Organic matter (%) | 25.8 ± 1.2 |
| P2O5 Olsen (%) | 319.2 ± 8.7 |
| K2O (mg/kg) | 1011.2 ± 19.7 |
| MgO (mg/kg) | 501.7 ± 8.4 |
| CaO (mg/kg) | 3940 ± 99.6 |
| Na2O (mg/kg) | 103 ± 2.2 |
| N (mg/kg) | 3674.2 ± 180.6 |
| C (g/kg) | 149.9 ± 7.1 |
| C/N ratio | 40.8 ± 0.9 |

In April 2024, pots with and without competition were divided into high and low canopy opening levels (Figure 1). One tunnel was set up in an open area of the experimental garden (high canopy opening), and another under the tree canopy (low canopy opening). Three Photosynthetically Active Radiation (PAR) detectors (tube of 30 cm length, ©SOLEMS DPAR/LEC1C#) were placed in each tunnel to assess transmitted irradiance. Three additional detectors were placed in open conditions, outside the experimental garden (*i.e*., under full light conditions). Measurements were recorded every minute for 24 h, in April 2024 on a clear-sky day. Light transmittance was computed as the ratio of irradiance in each opening condition to irradiance under full light conditions. Light transmittance was uniform under each tunnel, 37% under the high opening treatment, compared to only 10% under the low opening treatment. A similar difference was observed at the Saint-Christol-d’Albion study site between unthinned control plots and plots thinned at 75% of the initial basal area (Brasseur et al., 2025). Temperature and relative humidity were recorded throughout the experiment every 5 min using a HOBO data logger (Onset Computer Corporation, Bourne, MA, USA) in both tunnels (Table 2). Vapor pressure deficit (VPD in Pa) was calculated according to the function:

**Table 2.**
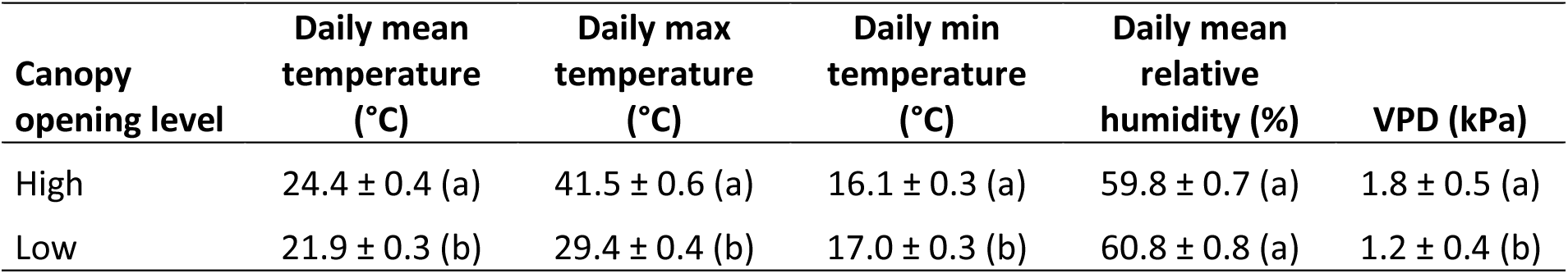
Means ± standard errors of daily mean, maximum, and minimum temperature (in °C), daily mean relative humidity values (in %) and daily VPD (Vapor Pressure Deficit in kPa) over the experimental period. Different letters indicate significant differences between canopy opening levels (Kruskal Wallis tests with *p*-value threshold = 0.05).

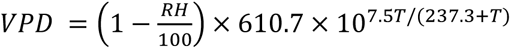 with *T* the temperature in °C and *RH* the relative humidity in % (Prévosto et al. 2020b).

For three months, from July 2024 to September 2024, half of the pots in each canopy opening treatment were subjected to reduced water supply to maintain them at 30–40% of field capacity, corresponding to a soil water deficit treatment, while the other half were maintained at 70–80% of field capacity (favorable soil water conditions) (Figure 1). Soil moisture was maintained using an automatic, controlled drip irrigation system and monitored by weighing the pots every 2 to 3 days. Our irrigation system compensates for differences in water consumption among the various treatment groups. We have chosen to focus on maintaining all pots at a certain percentage of field capacity, which results in differences in the amount of water supplied depending on the treatment group (due, for example, to differences in evapotranspiration between two irradiance treatments).

Seedlings were regularly monitored throughout the experiment. No visual symptoms of root rot, damping-off, or leaf pathogenic infections were observed during the experiment.

### 2.3. Festuca ovina measurements

In September 2024, *F. ovina* clumps were harvested, and the above- and belowground parts were separated. Roots were carefully washed over a sieve to remove soil particles without loss. The above- and belowground parts were then frozen, freeze-dried and weighed.

### 2.4. Quercus pubescens growth

*Quercus pubescens* seedlings size (principal stem height - from the root collar to the apical bud - in cm and diameter at the root collar in mm) was measured at the beginning of the experiment (October 2023), during the growing season before water stress was induced (July 2024), and at the end of the experiment (September 2024). We calculated the Relative Growth Rate (RGR) for height and diameter as:

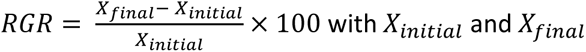 with *X_initial_* and *X_final_* the height or diameter at the beginning and at the end of the experiment, respectively.

The height/diameter ratio was calculated and the number of leaves was also recorded for each seedling.

In September 2024, *Q. pubescens* seedlings were harvested, and their root systems were carefully separated from *F. ovina* clumps. The root systems were then washed over a sieve to remove soil particles without losing any roots. Seedlings were separated into leaves, stems, primary roots (diameter > 2 mm) and fine roots (diameter < 2 mm) then frozen, freeze-dried and weighed.

### 2.5. Quercus pubescens physiological response and mycorrhization

In September 2024, we measured the chlorophyll index on 3 leaves per seedling, expressing the relative chlorophyll content per unit leaf area, using a SPAD-502 Plus Chl (Konica Minolta Optics, Japan). We also scanned every leaf of each seedling to measure their area and estimate the SLA (mm²/mg).

To quantitatively assess mycorrhizal colonization of *Q. pubescens* fine roots, we used Phospholipid fatty acid (PLFA) analysis following the method described in Biryol et al. (2024). We measured mycorrhizal biomass in µg per gram of dry fine root previously grounded with a ball mill (Retch) (collection and freeze-drying process described above). PLFA 18:2ω6,9 was used as a marker molecule for fungal biomass (Bååth & Anderson, 2003; Frostegård & Bååth, 1996; Klamer & Bååth, 2004; Wallander et al., 2013). This fungal biomarker is commonly used in soil microbial community studies but has also been found in plant tissues (Laczko et al., 2004; Olsson, 1999; Zelles, 1997). However, as described by Kaiser et al. (2010), most of the PLFAs measured in roots originate from ectomycorrhizal fungi growing inside the roots.

### 2.6. Soil physico-chemical analyses

Initial substrate samples were collected at the beginning of the experiment. At the end of the experiment in September 2024, soil samples were collected during seedling harvesting. All soil samples were dried (60°C for 48h in a drying oven) and sieved using a 2 mm sieve. Soil physico-chemical analyses were then carried out by Teyssier laboratory (Bordeaux, France, www.laboratoire-teyssier.com). Soil pH (measured in water and KCl), total organic matter and organic carbon (C, determination after dry combustion), total nitrogen (N, Kjeldahl method), phosphorous (P as P_2_O_5_, Olsen method), potassium (K as K_2_O), magnesium (Mg as MgO), calcium (Ca as CaO), sodium (Na as Na_2_O) and cation exchange capacity (CEC) were analyzed for each soil samples (methods are specified in Supplementary material Table S1).

### 2.7. Statistical analyses

All statistical analyses were performed using the R software 2024.12.1 +563 (R version 4.3.1, Foundation for Statistical Computing, Vienna, Austria).

The effects of canopy opening and soil water deficit, as well as their two-ways interaction on *F. ovina* belowground and aboveground biomass were tested using linear models and type III analysis of variance.

The effects of competition, canopy opening, soil water deficit, their two-way interactions and the three-way interactions on *Q. pubescens* morphological and physiological traits (height and diameter at the end of the experiment, height/diameter ratio, aboveground biomass, belowground biomass, aboveground/belowground biomass ratio, number of leaves, SLA, mycorrhization) and on each soil physico-chemical parameter were tested using linear or generalized linear models and type III ANOVA depending on the evaluated trait.

The effects of time, competition, canopy opening, soil water deficit and their two-way interactions on *Q. pubescens* growth parameters (height, diameter and height/diameter ratio) were tested using linear models (with log(x) transformation) and type III analysis of variance (ANOVA).

The normality of residuals was assessed through histogram inspection, Shapiro-Wilk test, and Q-Q plot visualization. Transformed data were used when needed to ensure residuals normality for linear models. Quasi-poisson family models were used in case of overdispersion of Poisson family models. All models, data transformation performed and *p*-value of Shapiro-Wilk tests and R² associated are summarized in Supplementary materials (Supplementary Tables S2, S3, S4, S5. B and S7). If model diagnostics indicated the presence of one influential observation (studentized residual > 3 and high Cook’s distance), the data point was considered an outlier and was excluded from the analysis to meet model assumptions of normality and homoscedasticity (this model diagnostics indicated the presence of only one influential observation, Supplementary Tables S3). Model predictions were plotted according to the different modalities with the 95% confidence intervals calculated as: *ŷ* ± *t*_(1−∝/2,*df*)_ × *SE*(*ŷ*) where *ŷ* is the predicted value, *t*_(1−∝/2,*df*)_ is the critical value of Student’s *t* distribution according to the degree of freedom and *SE*(*ŷ*) is the standard error of the predicted value.

Residual’s normality was not verified for chlorophyll index data. We therefore used non-parametric tests. We performed a Kruskal-Wallis test with post hoc Dunn test (with false discovery rate correction) to determine which modalities differ significantly from the others.

To assess the effect of competition, canopy opening and soil water deficit on soil physico-chemical profiles, we used a Principal Component Analysis (PCA) on standardized data. We performed a permutational multivariate analysis of variance (PERMANOVA) on the PCA individual coordinates using the adonis function of the Vegan package.

## 3. Results

### Festuca ovina biomass

Above- and belowground biomass of *F. ovina* were respectively 1.6 and 2.7-fold higher under high compared to low canopy opening. Aboveground biomass was negatively affected by soil water deficit only under high canopy opening (-28.6%), while the belowground biomass was not influenced by soil water availability (Table 3, Figure 2. A).

**Figure 2.**
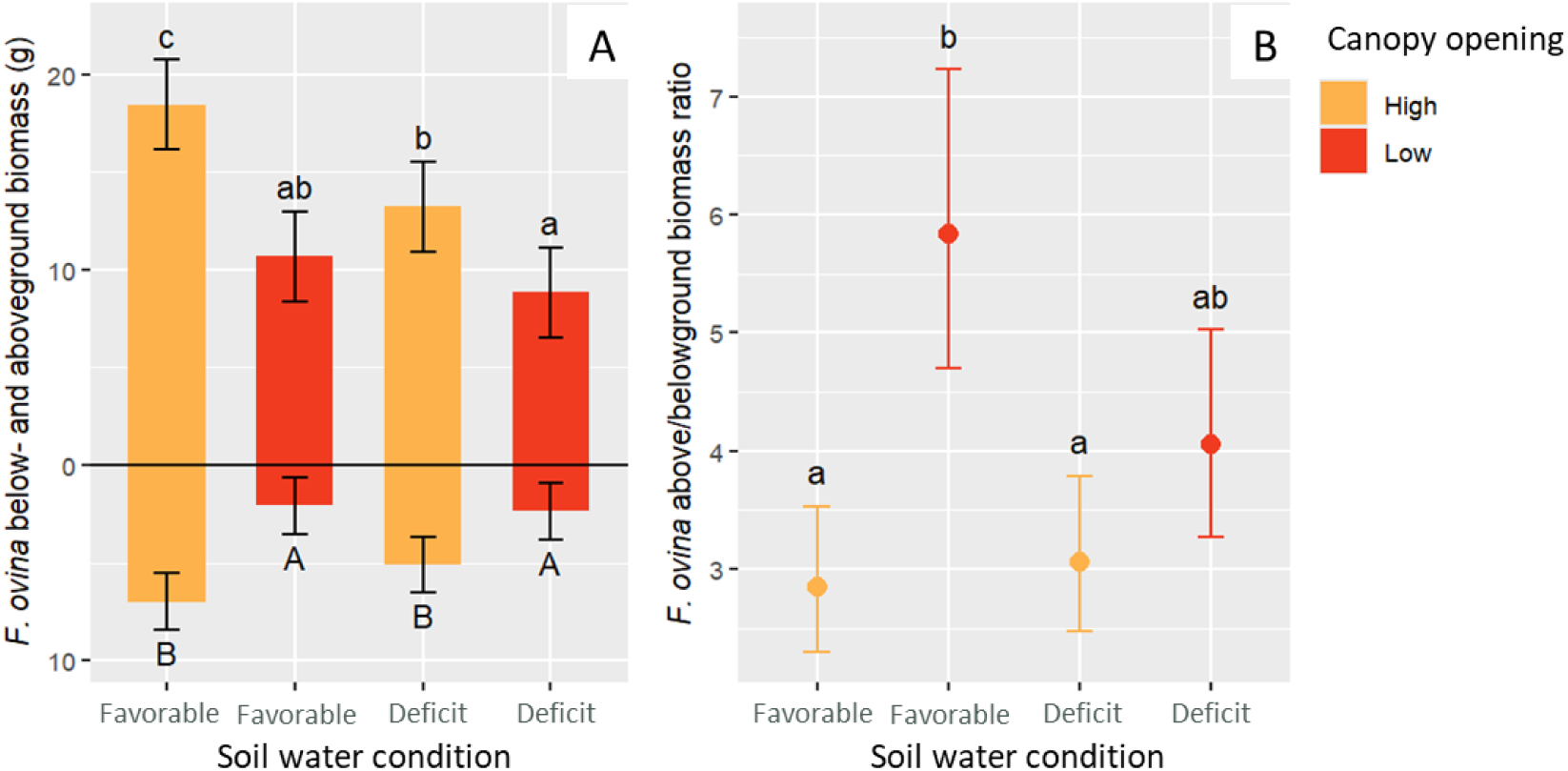
*Festuca ovina* above- and belowground biomass in g (A) and above/belowground ratio (B). Bars represent means ± standard error, n=12. Letters indicate significant differences between treatments based on post hoc Tukey tests (p-value < 0.05). (A) Uppercase letters below the bars indicate significant differences in belowground biomass; lowercase letters above the bars refer to aboveground biomass.

**Table 3.** Results of type III ANOVA and associated P values testing for the effects of canopy opening (O), soil water conditions (W) and their interaction on *F. ovina* traits. (Notes: * for P < 0.5, ** for P < 0.01 and *** for P < 0.005)

|  | Canopy Opening (O) | Soil water conditions (W) | O × W |
|---|---|---|---|
| *F. ovina a*boveground biomass (g) | **28.11\*\*\*** | **9.49\*\*** | 2.18<sup>*ns*</sup> |
| *F. ovina below*ground biomass (g) | **135.96\*\*\*** | **28.97\*\*\*** | 1.29<sup>*ns*</sup> |
| Aboveground/belowground ratio | **624.96\*\*\*** | **22.01\*\*\*** | 1.88<sup>*ns*</sup> |

### Quercus pubescens growth

First, a positive correlation was observed between time and *Q. pubescens* seedling growth with an average height relative growth rate (RGR) of 124.8% and an average diameter RGR of 64.6% across all treatments between the beginning and the end of the experiment (Table S5 and Figure S1 in Supplementary material).

In the absence of competition, *Q. pubescens* seedling height and diameter at the end of the experiment were 108.2% and 21.1% higher under high than under low canopy opening (Table 4 and Figure 3). Under high canopy opening, a negative effect of soil water deficit was only observed on height (-46.6%) (Table 4 and Figure 3). Under low canopy opening, soil water availability did not influence height or diameter.

**Figure 3.**
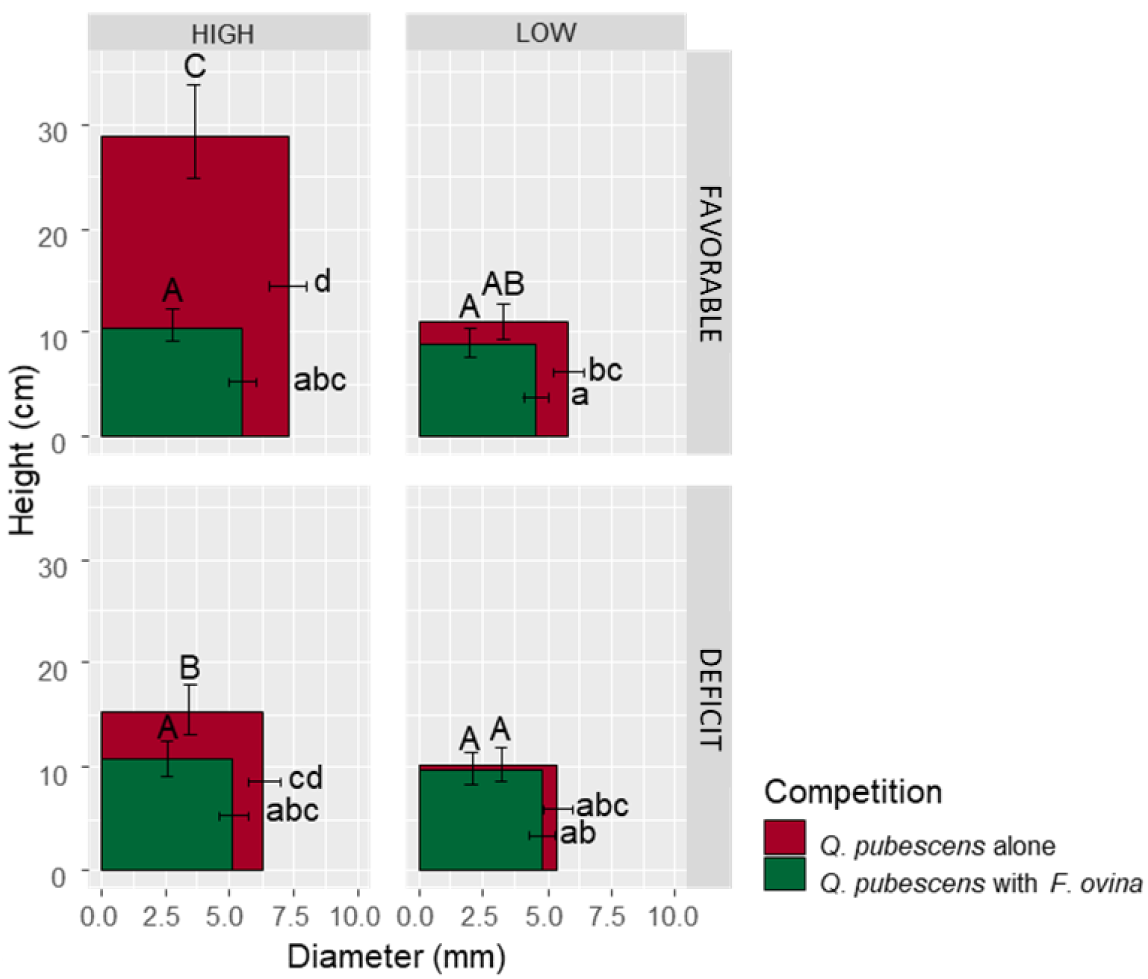
Effects of competition, canopy opening, soil water conditions and their interactions on *Q. pubescens* growth in height and diameter. Predicted values are obtained from linear and generalized linear models with Anova (type III). The letters indicate significant differences (uppercase for height and lowercase for diameter) based on Tukey’s post hoc comparisons (p-value < 0.05) within each combination.

**Table 4.** Results of type III ANOVA and associated P values testing for the effects of competition (C), canopy opening (O), soil water conditions (W) and their interaction on *Q. pubescens* growth parameters at the end of the experiment (height, diameter and height/diameter ratio). (Notes: * for P < 0.5, ** for P < 0.01 and *** for P < 0.005).

|  | Competition (C) | Canopy Opening (O) | Soil water conditions (W) | C × O | C × H | O × H | C × O × H |
|---|---|---|---|---|---|---|---|
| Height (cm) | **52.8\*\*\*** | **56.1\*\*\*** | **7.7\*\*** | **25.1\*\*\*** | **13.4\*\*\*** | **8.5\*\*** | **4.6\*** |
| Diameter (mm) | **36.5\*\*\*** | **20.2\*\*\*** | 2.7<sup>*ns*</sup> | 0.8<sup>*ns*</sup> | 2.0<sup>*ns*</sup> | 2.0<sup>*ns*</sup> | 0.2<sup>*ns*</sup> |
| Height/diameter | **12.0\*\*\*** | **20.8\*\*\*** | 2.8<sup>*ns*</sup> | **19.3\*\*\*** | **7.4\*\*** | **4.3\*** | **6.1\*** |

Under high canopy opening level and favorable soil water availability, the presence of *F. ovina* significantly reduced *Q. pubescens* growth in height (-63.3%) and diameter (-24.4%) at the end of the experiment (Table 4 and Figure 3). The height RGR between the beginning and the end of the experiment was 395.6% for *Q. pubescens* alone, while it reached only 93.8% in presence of *F. ovina* (Table S5 and Figure S1. A in Supplementary material). A similar trend was noted for the diameter but with less intense effect of *F. ovina*: a 112.1% RGR without competition compared to a 71.3% RGR in presence of *F. ovina* (Table S5 and Figure S1. B in Supplementary material).

Under low canopy opening, no effect of competition was detected on height growth. However, low canopy opening attenuated the effect of competition on diameter growth (-21.0% *versus* -24.4% (Table 4 and Figure 3)).

Soil water availability modulated the effect of competition on *Q. pubescens* growth (significant interaction between Competition × Soil water deficit for height growth, Table 4). Under soil water deficit, the negative effect of competition on growth was strongly attenuated for height (-32.2% *versus* -63.3%) and alleviated for diameter (Figure 3).

Low canopy opening and soil water deficit alleviated the effect of competition on both height and diameter growth (Figure 3).

### Quercus pubescens biomass

In the absence of competition, both above- and belowground biomass of *Q. pubescens* seedling increased under high canopy opening (Table 5 and Figure 4. A). Under favorable soil water conditions, above- and belowground biomass was 424.6% and 167.5% higher under high compared to low canopy opening, respectively.

**Figure 4.**
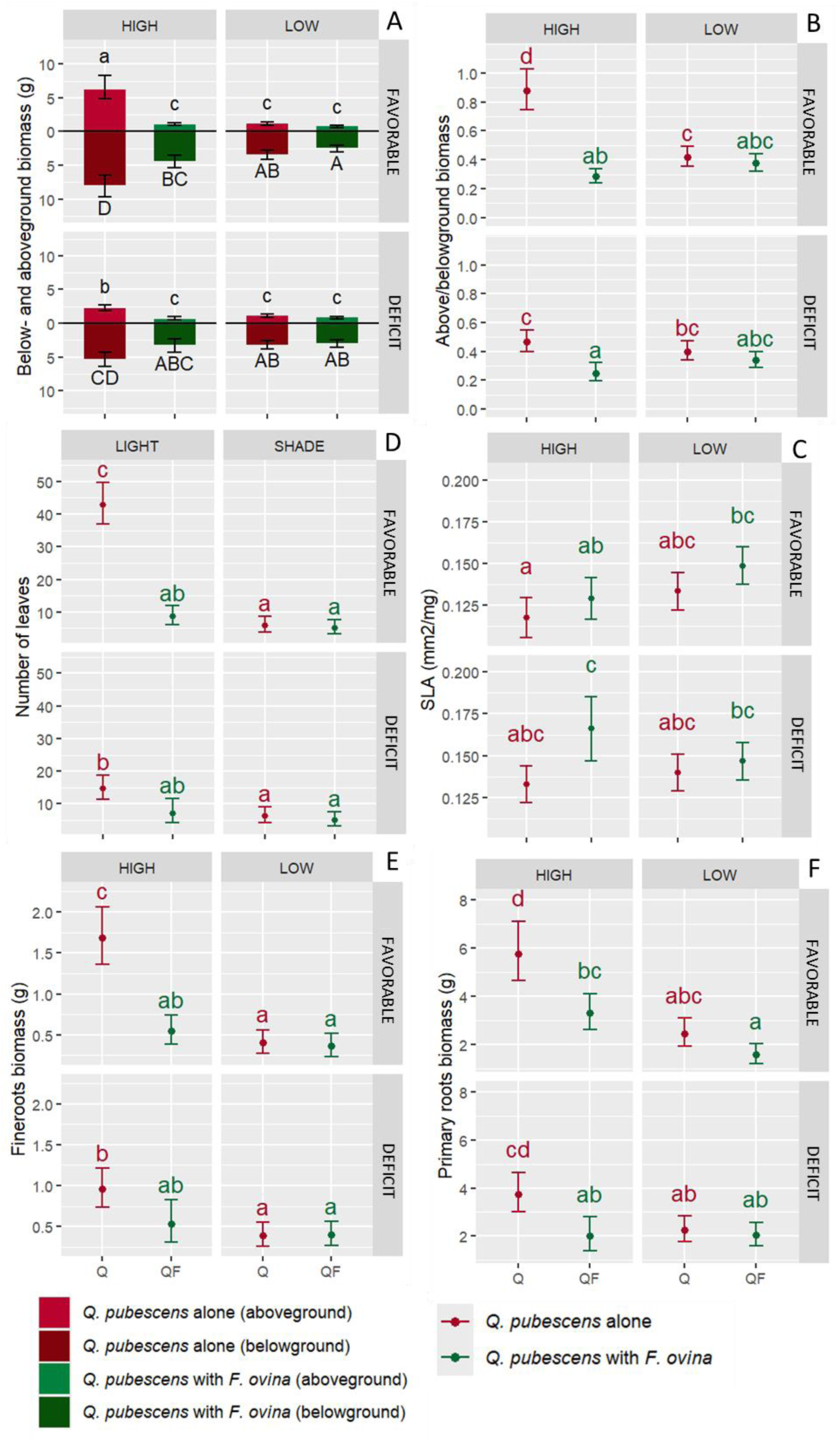
Effects of competition, canopy opening, soil water conditions and their interactions on *Q. pubescens* aboveground and underground biomass per individual (A), aboveground/underground biomass ratio (B), SLA (C), Number of leaves per individual (D), Fine roots biomass per individual (E) and Primary roots biomass per individual (F). Predicted values are obtained from linear and generalized linear models (Table 5) with Anova (type III).

**Table 5.** Results of type III ANOVA and associated P values testing for the effects of competition (C), canopy opening (O), soil water conditions (W) and their interaction on *Q. pubescens* morphological and physiological traits. (Notes: * for P < 0.5, ** for P < 0.01 and *** for P < 0.005)

|  | Competition (C) | Canopy Opening (O) | Soil water conditions (W) | C × O | C × W | O × W | C × O × W |
|---|---|---|---|---|---|---|---|
| Aboveground biomass (g) | **131.3\*\*\*** | **64.5\*\*\*** | **20.4\*\*\*** | **48.1\*\*\*** | **5.6\*** | **21.6\*\*\*** | 1.2<sup>*ns*</sup> |
| Underground biomass (g) | **23.2\*\*\*** | **43.7\*\*\*** | **4.1\*** | **5.6\*** | 1.4<sup>*ns*</sup> | **7.9\*\*** | 0.3<sup>*ns*</sup> |
| Aboveground/underground | **65.5\*\*\*** | 1.6<sup>*ns*</sup> | **13.2\*\*\*** | **34.5\*\*\*** | 3.1<sup>*ns*</sup> | **6.2\*** | **5.0\*** |
| Fine roots biomass (g) | **19.5\*\*\*** | **46.1\*\*\*** | 2.3<sup>*ns*</sup> | **17.6\*\*\*** | 3.2<sup>*ns*</sup> | 3.2<sup>*ns*</sup> | 1.8<sup>*ns*</sup> |
| Primary roots biomass (g) | **25.2\*\*\*** | **37.2\*\*\*** | **5.4\*** | **4.2\*** | 0.7<sup>*ns*</sup> | **10.1\*\*** | 1.0<sup>*ns*</sup> |
| Leaf biomass (g) | **173.3\*\*\*** | **95.8\*\*\*** | **43.2\*\*\*** | **89.2\*\*\*** | **15.3\*\*\*** | **36.3\*\*\*** | **11.1\*\*** |
| Stem biomass (g) | **84.8\*\*\*** | **46.9\*\*\*** | **14.0\*\*\*** | **28.4\*\*\*** | **8.2\*\*** | **19.1\*\*\*** | 2.2<sup>*ns*</sup> |
| SLA (mm<sup>2</sup>/mg) | **14.0\*\*** | 1.7<sup>*ns*</sup> | **10.0\*\*** | 1.6<sup>*ns*</sup> | 0.5<sup>*ns*</sup> | **7.1\*** | 2.9<sup>*ns*</sup> |
| Leaf surface (mm<sup>2</sup>) | 2.9<sup>*ns*</sup> | **5.3\*** | 0.1<sup>*ns*</sup> | 1.5<sup>*ns*</sup> | 0.04<sup>*ns*</sup> | 1.3<sup>*ns*</sup> | 0.1<sup>*ns*</sup> |
| Number of leaves | **27.1\*\*\*** | **46.0\*\*\*** | **5.7\*** | **13.4\*\*\*** | 2.0<sup>*ns*</sup> | **6.1\*** | 3.1<sup>*ns*</sup> |
| Fungal abundance (µg/g) | **32.5\*\*\*** | 3.9<sup>*ns*</sup> | **6.6\*** | **12.3\*\*\*** | 0.7<sup>*ns*</sup> | 0.3<sup>*ns*</sup> | 0.03<sup>*ns*</sup> |
| SQI (Seegling Quality Index) | **11.7\*\*** | **20.7\*\*\*** | 2.9<sup>*ns*</sup> | 0.4<sup>*ns*</sup> | 0.003<sup>*ns*</sup> | 3.3<sup>*ns*</sup> | 3.4<sup>*ns*</sup> |

A negative effect of soil water deficit on the aboveground biomass was observed, but only under high canopy opening (-67.0%) (significant interaction between canopy opening × soil water condition, Table 5). Belowground biomass was not influenced by soil water availability (Figure 4. A).

The same response was observed for most seedling compartments (leaves, stems and fine roots biomass) as well as for the number of leaves and the ratio between above- and belowground biomass (Table 5, Figure 4. B, D, E and Figure S2 in Supplementary material), in response to canopy opening and soil water deficit, while primary roots biomass followed a similar trend to that of total belowground biomass (Table 5, Figure 4. F).

Under high canopy opening and under favorable soil water availability, the presence of *F. ovina* significantly reduced *Q. pubescens* aboveground (-84.2%) and belowground (-54.0%) biomass (Table 5 and Figure 4. A). A similar response pattern to competition was observed for all seedling biomass compartments (leaves, stems, fine roots biomass), for the number of leaves and for the aboveground/belowground ratio (Table 5, Figure 4. B, D, E and Figure S2. A, C in Supplementary material).

The negative effect of competition on aboveground and belowground biomass disappeared under low canopy opening (significant interaction between competition × canopy opening, Table 5, Figure 4. A). Soil water deficit attenuated the negative effect of competition on aboveground biomass (-69.5% *versus* -84.2%) and alleviated the effect of competition on belowground biomass (significant interaction between competition × soil water deficit, Table 5, Figure 4. A).

Low canopy opening and soil water deficit limited the effect of competition on both aboveground and belowground biomass (Figure 4. A).

A similar response pattern to that described for aboveground biomass was observed for abiotic control of competition impact on *Q. pubescens* leaf, stem and primary roots biomasses, as well as on the ratio between aboveground and belowground biomass. Fine roots biomass and leaf number followed the same response pattern of belowground biomass (Table 5, Figure 4. B, D, E, F and Figure S2 in Supplementary material).

### Quercus pubescens leaf traits

*Quercus pubescens* leaf area was not affected by canopy opening, soil water deficit or competition (Table 5, and Figure S2 in Supplementary material). However, the SLA of seedlings grown under high canopy opening, without *F. ovina* and under favorable soil water availability were smaller than that of seedlings grown with *F. ovina* under at least one stressful abiotic condition (low light, water deficit, or both) (Table 5 and Figure 4. C).

In the absence of competition, *Q. pubescens* chlorophyll index was 6.6% lower under low than under high canopy opening. We observed a positive effect of soil water deficit on the chlorophyll index (+5.3%) but only at a high canopy opening level (Figure 5). Under low canopy opening, soil water deficit did not influence the chlorophyll index. Under high canopy opening and under favorable soil water availability, the presence of *F. ovina* reduced the *Q. pubescens* chlorophyll index (-33.4%), but this negative effect of competition was attenuated under low canopy opening level (-12.3%) (Figure 5). Soil water deficit did not influence the effect of competition on chlorophyll index.

**Figure 5.**
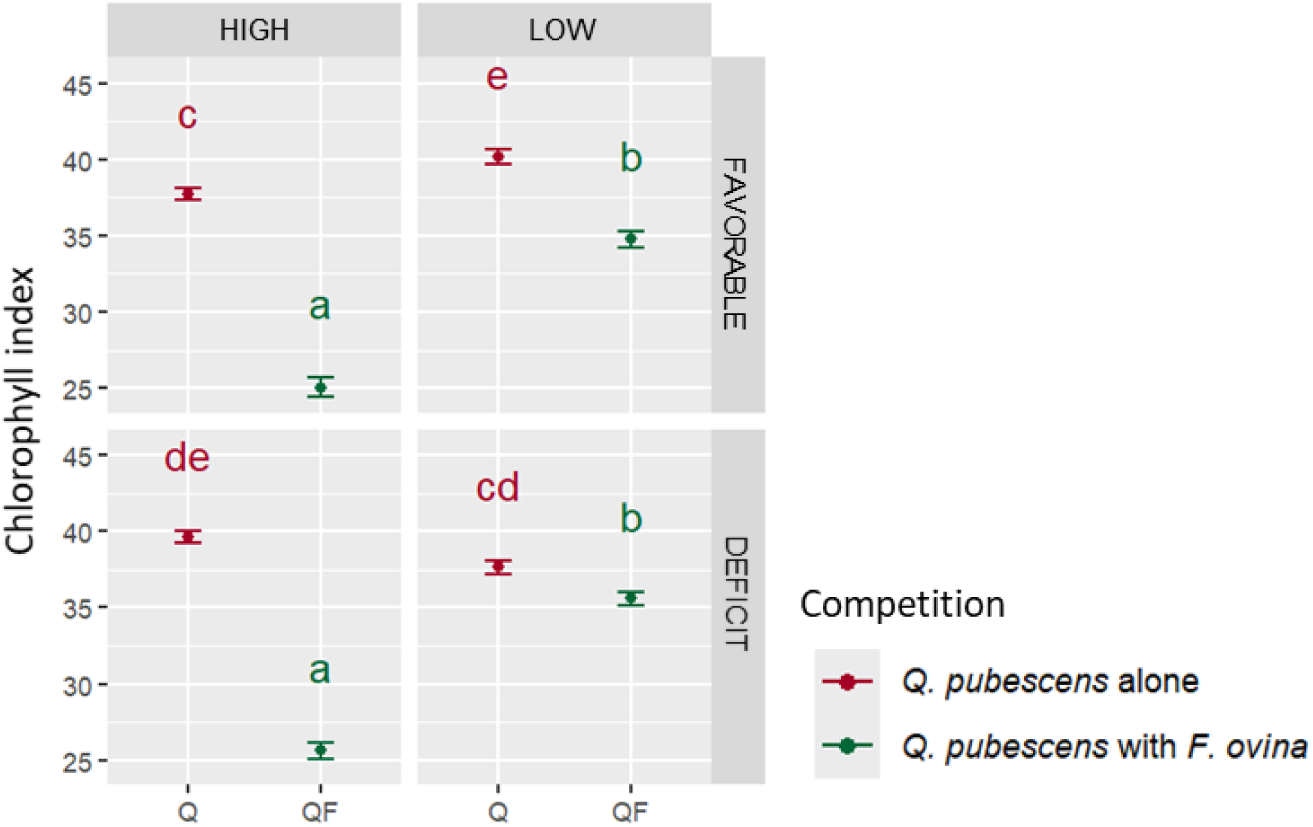
Effects of competition, canopy opening, soil water conditions and their interactions on *Q. pubescens* chlorophyll index (mean ± se). Kruskal-Wallis p-value < 2.2e^-16^ and Dunn test results are shown with letters.

### Mycorrhization

In the absence of competition, fungal abundance associated with *Q. pubescens* roots was not affected by canopy opening levels nor soil water deficit (Figure 6).

**Figure 6.**
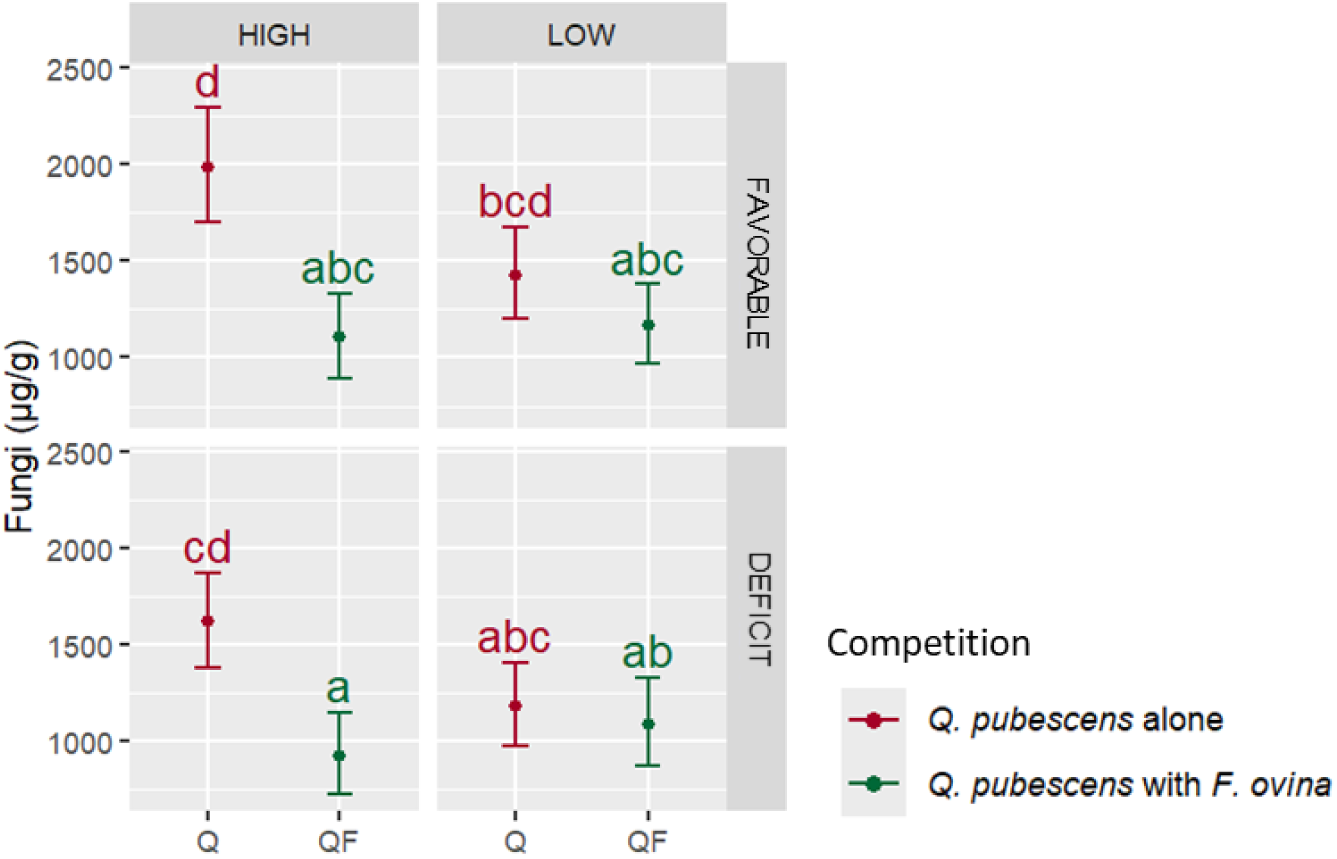
Effects of competition, canopy opening, soil water conditions and their interactions on the fungal abundance associated to *Q. pubescens* roots (µg/g of dry fine roots). Predicted values are obtained from a linear model (Table 5) with Anova (type III).

Under high canopy opening, and under favorable soil water availability, the presence of *F.* ovina induced a marked decrease in fungal abundance associated to *Q. pubescens* roots (-44.6%) (Figure 6). Under soil water deficit, this negative effect of competition on fugal abundance was still significant (-42.7%). Under low canopy opening, the presence of *F. ovina* did not influence the fugal abundance (Figure 6).

### Soil physico-chemical parameters

Principal component analysis (PCA) of soil physico-chemical parameters showed that the first PCA axis explaining 44.5% variation loaded with high scores of CEC, pH (water and KCl) and Ca, Mg and Na cations concentrations) (Figure 7. A) The soil profiles corresponding to the presence or absence of *F. ovina* were distinguished along this first axis of the PCA (*P*_permanova_ < 0.001, Figure 7. B). The presence of *F. ovina* positively influenced the cation-exchange capacity (CEC), regardless of abiotic conditions (Table S6 and Figure S3 in Supplementary material). Under favorable soil water availability, the presence of *F. ovina* positively influenced pH, and the concentration in MgO, CaO and Na_2_O (Table S6 and Figure S3 in Supplementary material).

**Figure 7.**
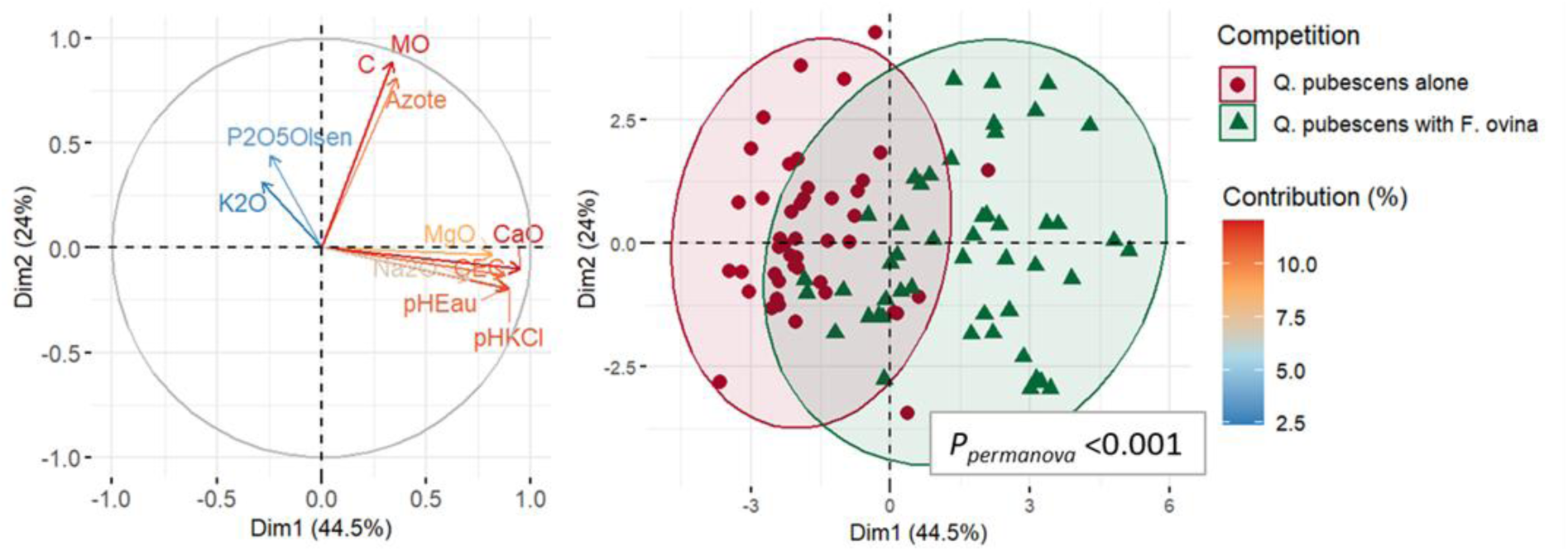
Principal component analysis (PCA) from the soil physico-chemical parameters according to competition levels and main explanatory variables of PC1 and PC2.

## 4. Discussion

### Effects of abiotic factors on Q. pubescens seedlings

Our experiment highlighted the positive effect of canopy opening on *Q. pubescens* growth as predicted by Hypothesis 1. Canopy opening leads to an increase in both temperature and light availability (Mölder et al., 2019), the latter being the primary driver of growth stimulation (Espelta et al., 1995). *Quercus pubescens* is considered to be a light-demanding species (Contran et al., 2013; Pasta et al., 2016), and light availability is particularly important for oak regeneration (Mölder et al., 2019). Consistent with other studies on deciduous oak species, our results demonstrate an increase in both aboveground and belowground seedling biomass with increasing light availability (Sevillano et al., 2016; Wang & Bauerle, 2006; Welander & Ottosson, 1998). We observed that *Q. pubescens* seedlings tend to allocate more biomass to their aboveground than their belowground parts under high light availability, indicating a preferential investment in photosynthetic organs under optimal conditions (Kozlowski & Pallardy, 2002). Limited light availability in the understory of mature forests is widely recognized as a major constraint on the survival and growth of oak seedlings (Dey et al., 2012), a pattern that our results also confirm by highlighting limited growth under low light conditions. This limitation explains the traditional reliance on clear-cutting to stimulate regeneration, though this primarily favors vegetatively reproduction (*i.e*., for oaks, sprouts over seedlings from acorns) (Brasseur et al. 2025) and competition by the ground vegetation (see discussion below). On the other hand, consistent with the findings of Kwak et al. (2011) on *Q. suber*, we observed that chlorophyll content was higher under low compared to high light conditions. This suggests an efficient shade-acclimation response in *Q. pubescens* through enhanced investment in light-harvesting pigments under limited light availability, a strategy also observed under drought stress (Gallé et al., 2007; Laoué, Gea-Izquierdo, et al., 2024; Laoué, Havaux, et al., 2024). We also observed a lack of difference in the abundance of root-associated fungi between high and low light conditions. This contrasts with studies on other species such as that of Shi et al. (2014) or Shukla et al. (2009) who reported a positive effect of light on root mycorrhizal colonization, suggesting a species-specific or context-dependent response.

Our study demonstrates, as expected, that soil water deficit negatively affects *Q. pubescens* growth (in line with Hypothesis 1), particularly in aboveground parts, leading to decreases in height, leaf number and leaf and stem biomass. Despite the slowdown in growth, *Q. pubescens* – a species known for its moderate drought tolerance (Gallé et al., 2007; Haldimann et al., 2008; Saunier et al., 2022) - exhibited several strategies to cope with soil water deficit. First, seedlings showed a shift in biomass allocation toward belowground parts, with an investment concentrated in primary roots, which did not appear to be affected by the soil water deficit. This pattern suggests an adaptative strategy which results in increasing the volume of soil exploited for water supply, as already observed by Di lorio et al. (2011). We further observed a reduction in fine roots biomass that could indicate an increased vulnerability of fine roots to desiccation but also indicate a shedding strategy (Di lorio et al., 2011): Under water stress, the maintenance costs for *Q. pubescens* seedlings are reduced, and the role of thicker roots as temporary water storage organs is enhanced (Eissenstat & Yanai, 1997). Conversely, water stress caused an increase in leaf chlorophyll content, indicating that *Q. pubescens* investment in light-harvesting pigments are stimulated under limiting conditions, a response that we also observed when *Q. pubescens* is under low light availability. Finally, soil water deficit did not affect the abundance of root-associated fungi. While water stress can variably influence mycorrhizal colonization (positively or negatively), maintaining an effective mycorrhizal symbiosis is essential for plant drought tolerance as it helps increase water uptake by the plant.

Finally, soil water deficit did not affect fungal abundance when expressed per unit of root mass. This result should be interpreted considering the observed decrease in fine root biomass. Although fungal abundance at the root level was not influenced by soil water deficit, the overall reduction in fine root biomass was not counterbalanced by an increase in mycorrhizal colonization. However, effective mycorrhizal symbiosis is essential for plant drought tolerance, as it can improve water and nutrient uptake and mitigate low water stress (Augé, 2001; Domínguez Núñez et al., 2009; Mohan et al., 2014). Augé (2001) reported that drought can have contrasting effects on root colonization but that it more frequently leads to increased colonization, particularly under field conditions. Since the present study was conducted as a pot experiment, future field-based research will be vital to clarify whether these patterns hold true in more complex ecosystems.

For most of the morphological traits measured, we observed a strong interaction in their response to canopy opening (*i.e*., light availability) and soil water deficit. According to the facilitation hypothesis, under drought conditions, the negative effects of light limitation may be outweighed by its benefits for plant water status (Callaway, 1995; Sack, 2004; Valladares et al., 2016). Our study supports this hypothesis, as we observed a limited effect of soil water deficit on most *Q. pubescens* seedlings traits under low light conditions. As discussed by Maestre et al. (2003), Quero et al. (2006) or Sánchez-Gómez et al. (2006), the facilitating role of shade during periods of drought could be involved in the regeneration process in Mediterranean environments. This has direct implications for forest management practices that modulate forest canopy cover, such as thinning. However, it should be noted that, under combined shade and soil water deficit, we observed that *Q. pubescens* seedlings failed to enhance photosynthesis, in contrast to single stress conditions. Similar findings for *Q. suber* suggest that shade can impair the development of key drought-tolerance mechanisms, including osmotic adjustment and water loss regulation (Aranda et al., 2005). Our results also indicate that only the combined effect of shade and soil water deficit significantly limits root colonization by mycorrhizae. It is important to emphasize here that, for feasibility reasons, the experiment was conducted in Marseille, i.e., under warmer climatic conditions than those prevailing in the supramediterranean belt from which the *Q. pubescens* acorns originated. These warmer conditions may have had an influence on our results. For example, soil water deficit and stress effects may have been exacerbated by high temperatures (Haldimann et al., 2008). We highlight here the need to conduct additional complementary experiments, particularly under natural field conditions, in order to further support and complete the results presented in this study.

### Effects of competition on Q. pubescens seedlings

First, our results highlight the effect of competition with a Poaceae species on *Q. pubescens* seedling growth, confirming the importance of considering the herbaceous stratum in the regeneration process (Thrippleton et al., 2016), and in accordance with Hypothesis 2. Indeed, in the presence of *F. ovina*, we observed reduced growth (height, diameter, above- and belowground biomass) as well as a lower above-/belowground biomass ratio compared to *Q. pubescens* seedlings developed without *F. ovina.* This latter observation indicates a higher investment in root biomass which results in greater resistance to competition for water and nutrient resources, which is in line with the findings of Kunstler et al. (2006). Our results are also consistent with those of Tonioli et al. (2001), who demonstrated in a field experiment that the surrounding vegetation, even though it plays a facilitating role during emergence, competes with young *Q. pubescens* seedlings for their growth. Another pot experiments, described by Coll et al. (2004), on *Fagus sylvatica* demonstrated the strong competitiveness of Poaceae species, particularly for water resources, thanks to their extensive root systems. Here, we also observed that the presence of *F. ovina* exerted a negative impact on the belowground biomass of *Q. pubescens*, affecting both fine and primary roots, but also inducing a reduction in chlorophyll index. In absence of *F. ovina*, oak seedlings displayed adaptative strategies to cope with soil water deficit, such as an increase in chlorophyll index (see discussion above). In contrast, no such compensatory responses were detected under competition in favorable soil water conditions, suggesting the stronger constraining effect of competition. Similarly, we observed a significant reduction in the abundance of root-associated fungi in the presence of *F. ovina*, an effect that was not found under soil water deficit or low light availability. Fernandez et al. (2023) studied the ectomycorrhization of *Q. petraea* seedlings growing with another Poaceae species, *Molinia caerulea,* and observed a reduced mycorrhizal association rate with oak roots. These inhibition patterns could be explained by allelopathic interactions, as discussed by Fernandez et al. (2023). Indeed, *F. ovina,* and more generally *Festuca* species, are known for their allelopathic potentialities (Bertin et al., 2007; de Bertoldi et al., 2012; Lipińska et al., 2013). Further experiments analyzing root exudates would provide valuable insights into the effects of *F. ovina* on *Q. pubescens* and its mycorrhization, by allowing us to distinguish between competitive effects and chemical interactions.

Furthermore, we observed that the presence of *F. ovina* also affects soil physicochemical characteristics. Our results highlight an increase in soil pH, cation exchange capacity (CEC), and the concentration of certain exchangeable cations (MgO, CaO, Na₂O) in the presence of *F. ovina*. These results are consistent with previous studies indicating that grasses can alter soil chemistry (Koukoura et al., 2003; Wyszkowska et al., 2022), thereby influencing the growth of woody plants. Indeed, a change in soil chemistry can affect soil structure and the availability of nutrients for the plants. Our results suggest that the presence of *F. ovina* is associated with improved soil fertility (particularly due to higher CEC and pH). However, this benefit does not appear sufficient to offset the negative effects of the presence of *F. ovina* on *Q. pubescens* growth. Further research would be needed to clarify the underlying mechanisms of competition.

It should be emphasized here that *F. ovina* was used as the competing species primarily for reasons related to the feasibility of conducting the experiment under semi-controlled conditions. In this context, the species should therefore be regarded as a model species. It would be valuable to extend the present analysis to experiments involving, for example, *F. rubra*, which, although sharing several characteristics typical of the genus *Festuca*, may also differ in its autecology, particularly through a lower tolerance to relatively dry soils (Rameau et al., 1989; Tichý et al., 2023). Nevertheless, our results confirm, as discussed for example by Kohler et al. (2020), that large canopy openings can constrain the regeneration of target species by increasing light availability and promoting the development of herbaceous species (Floret et al., 1992; Palviainen et al., 2005).

Similarly, it would be valuable to extend these findings to field conditions by focusing on other stages of acorn and seedling development. For instance, in natural environments, the influence of the herbaceous layer is evident from an early stage, affecting acorn desiccation. Dense herbaceous vegetation can create a mechanical barrier that prevents acorns from reaching the soil surface, thereby increasing the risk of desiccation (Löf et al., 2019). Conversely, once the herbaceous layer is established, particularly under open canopy conditions, it can mitigate desiccation by buffering microclimatic extremes and maintaining higher moisture near the soil surface (Tonioli et al., 2001).

### Interactive effects of abiotic conditions and competition on Q. pubescens seedlings

Our results, as expected in Hypothesis 3, also emphasize the interactions between biotic and abiotic factors, as we observed that the effects of competition depend on both canopy opening level and soil water conditions. Firstly, canopy opening, and therefore light availability, influences the competition between *F. ovina* and *Q. pubescens*. As expected for a Poaceae species (Xu et al., 2025), *F. ovina* above- and belowground biomass declined under low canopy opening, reflecting its sensitivity to light limitation. Furthermore, under low light conditions, the presence of *F. ovina* only marginally affected *Q. pubescens* seedling growth. In other words, as observed by Vernay et al. (2019) in the case of competition between *Q. petraea* and *Molinia caerulea*, competition interactions are amplified by light availability. Secondly, soil water deficit also influences the competition between *F. ovina* and *Q. pubescens*. At high canopy opening level, we observed a negative effect of soil water deficit on *F. ovina* aboveground biomass but not on its belowground biomass. This result corroborates previous observations indicating that the *Festuca* genus exhibits high drought tolerance, attributed to its fibrous rhizomes and extensive root system (Gibson & Newman, 2001; Yamada, 2011). Since water stress has a weaker impact on the growth of *F. ovina* than light limitation, competition between *F. ovina* and *Q. pubescens* is also less constrained by soil water deficit than by low light availability. Indeed, under water stress, the presence of *F. ovina* still negatively influenced *Q. pubescens* growth (*e.g.*, height, aboveground biomass or primary roots biomass). These results further highlight that intensive canopy opening can hinder regeneration. By increasing light availability, these openings intensify competitive interactions and exacerbate water stress, which disadvantage oak seedlings compared to *F. ovina*. However, it should be noted that low light availability imposes an equally strong limitation on oak seedling growth than water stress combined with competition under high light availability, showing that both intensive and insufficient canopy opening levels exert significant constraints.

Finally, our study contributes to the broader discussion on forest management recommendations aiming at enhancing regeneration in a climate change context. First, we highlight the role of competition with Poaceae species, in our case with a *Festuca* species, in oak regeneration. The presence of Poaceae species not only increases competitive pressure for resources, but also modifies soil properties and belowground interactions. Furthermore, our results emphasize the key roles of light availability and soil water conditions on seedling development, whose levels are largely dependent on silvicultural operations (Mölder et al., 2019). We revealed a persistent light requirement for optimal seedling performance, indicating that regeneration success depends on achieving a balance between sufficient canopy opening to provide light and enough cover to mitigate water stress and minimize grass competition. It would be interesting to continue our research with field studies to avoid the biases introduced by pot experiments, but also over a longer period of time, during which individuals experience several dry seasons.

## Acknowledgement

The study was supported by the European commission through the project “Holistic management practices, modelling and monitoring for European forest soils” - HoliSoils (EU Horizon 2020 Grant Agreement No. 101000289) and by the French government under France 2030 program, as part of the Aix-Marseille Université - A*MIDEX Excellence Initiative (AMX-19-IET-012) and the ECCOREV Research Federation (FR 3098: Aix-Marseille Univ., CNRS, INRAE, IRSN, CEA, Univ. Toulon, Univ. Avignon, Univ. Nîmes). This work was also supported by the French National program EC2CO (Ecosphère Continentale et Côtière). This work is also part of the REGE-ADAPT project of the research program FORESTT and received government funding managed by the Agence Nationale de la Recherche under the France 2030 program, reference ANR-24-PEFO-0006. We thank all those who helped during the field campaigns conducted at the study site of Saint-Christol: the IMBE and RECOVER laboratories and the CRPF PACA. We acknowledge Matthieu Jurado for the technical help in the experimentation performed at the IMBE experimental garden. We are grateful to the interns, Rachid Hafsaouy, Candice Martinod and Victoire Tharreau for their help with data collection and experiment monitoring.

